# Adaptation of a fluoroquinolone-sensitive *Shigella sonnei* to norfloxacin exposure

**DOI:** 10.1101/2020.12.08.416974

**Authors:** Bao Chi Wong, Soffi Kei Kei Law, Muhammad Zarul Hanifah Md Zoqratt, Qasim Ayub, Hock Siew Tan

## Abstract

*Shigella* causes shigellosis that requires antibiotic treatment in severe cases. Sub-lethal antibiotic concentrations can promote resistance, but their effect on antibiotic-sensitive bacteria before resistance development is unknown. This study investigated the effects of sublethal norfloxacin (NOR) challenges on a norfloxacin-sensitive strain, *Shigella sonnei* UKMCC1015. Firstly, the whole genome of *S. sonnei* UKMCC1015 was assembled, and 45 antimicrobial resistance (AMR) genes were identified. However, transcriptomic analysis showed that low NOR levels do not change the expression of AMR genes and norfloxacin targets. Instead, multiple ribosomal protein genes were downregulated, which could be attributed to decreased ribosomal protein promoter activity, modulated by elevated ppGpp levels. This alarmone is involved in the bacterial stringent response during environmental stress, and it is mainly produced from the ppGpp synthetase (*relA*). However, high levels of ppGpp (due to *relA* overexpression) are detrimental to the continued survival of *S. sonnei* during prolonged exposure to NOR. This study revealed that a norfloxacin-sensitive strain responds differently to sub-lethal NOR than commonly reported in resistant strains. Overexpression of *relA* (high ppGpp levels) may reduce the mutability of *S. sonnei* UKMCC1015 in response to NOR exposure and affect NOR tolerance.

## 2. Introduction

*Shigella* species is a common causal agent of episodic global diarrhoea, affecting mainly children younger than 5 years [1]. Due to the improving economic prosperity of many countries, *S. sonnei* has recently replaced *S. flexneri* as the dominant cause of bacillary dysentery globally [2]. The current population of *S. sonnei* descended from a common ancestor that led to distinct lineages and spread to other parts of the world, with strains in Asia that are mostly descendants of one lineage from Europe [3]. The etiological shift of shigellosis to *S. sonnei* has also been reported in Malaysia [4, 5]. Most studies on *Shigella* in Malaysia have focused on its antimicrobial susceptibility profiles, and there is a lack of data on its genetics and phylogeny.

The misuse and overuse of antibiotics have resulted in an increased concentration of antibiotics in the environment, exacerbating the antimicrobial resistance crisis [6]. Among all antimicrobial-resistant bacteria, fluoroquinolone-resistant *Shigella spp.* is the priority 3 (medium) category for new antibiotics research and development. Studies have shown low antibiotic concentrations might contribute to antimicrobial resistance [7–9]. Antibiotic concentrations higher than the minimum inhibitory concentration (MIC) lead to antimicrobial effects on susceptible cells. However, concentrations lower than the MIC (also known as sub-MIC) can act as a signal for diverse responses that ultimately induce tolerance [10]. As antibiotic pollution leads to sub-MIC levels of antibiotics in the environment, this condition can enrich pre-existing resistant variants and cause selective pressure to develop de novo resistance [11]. Thus, the effect of sub-MIC on antibiotic tolerance in sensitive bacterial strains needs to be well understood to counteract the inevitable presence of resistant strains.

Norfloxacin (NOR) is one of the first fluoroquinolones synthesised. Although it is not the preferred drug for treating shigellosis, its mechanism of action is similar to the recommended ciprofloxacin. All fluoroquinolones act by inhibiting DNA gyrase and topoisomerase IV. Fluoroquinolones bind at the enzyme-DNA interface, blocking ligation of the DNA and leading to lethal chromosomal breaks [12]. By inhibiting bacterial DNA synthesis, it will ultimately cause bacterial cell death. Exposure to sub-MIC fluoroquinolone can promote resistance via mutations in the DNA gyrase gene [13]. However, the transcriptional responses of a sensitive bacterium to low levels of fluoroquinolone before developing antimicrobial resistance are not well characterised.

This study determines the antibiotic susceptibility profile and characterises the *S. sonnei* strain UKMCC1015 from Malaysia. We present the first complete reference genome for *S. sonnei* from South East Asia. We also investigate the response of this fluoroquinolone-sensitive bacterium to a sub-lethal level of a second-generation fluoroquinolone NOR using RNA sequencing. The genes encoding the NOR target (DNA gyrase) are not differentially expressed. Instead, multiple ribosomal protein (r-protein) operons are significantly downregulated. We observe a slight decrease in several r-protein promoters and postulate that ppGpp may be involved in the r-protein inhibition upon short exposure to sub-lethal NOR. To investigate the role of ppGpp in long-term NOR exposure, we evolve UKMCC1015 *relA* mutants (with different intrinsic levels of ppGpp) in lethal doses of NOR using the adaptive laboratory evolution approach. Without *relA*, NOR resistance requires mutations in the DNA gyrase. Additionally, we observe that overexpression of *relA* confers moderate NOR resistance without mutating the DNA gyrase.

## 3. Materials and Methods

### Strain growth and culture conditions

Bacterial strains and plasmids used in this study are listed in Supplementary Table 1. UKMCC1015 strain is a clinical isolate purchased from the University Kebangsaan Malaysia Culture Collection (UKMCC). E. coli DH10β was used for general cloning purposes. The strain was cultured in Mueller-Hinton broth (Oxoid, UK) and incubated at 37°C with shaking at 200 rpm overnight. When necessary, 50 µg/mL of kanamycin (KAN), 100 µg/mL of ampicillin (AMP) and 15 µg/mL of chloramphenicol (CHL) were added to the culture media for selection. Total genomic DNA was extracted from the mid-log culture using the Wizard® Genomic DNA Purification Kit from Promega following the manufacturer’s protocol.

### Genome sequencing, assembly and annotation

We used a hybrid approach to obtain the complete genome, employing long-read Oxford Nanopore Technology and short-read Illumina MiSeq sequencing. Briefly, for long-read sequencing, DNA libraries were prepared using the Ligation Sequencing Kit (SQK-LSK109) according to the manufacturer’s protocol and sequenced on a MinION FLOW-MIN106 flowcell and MinION MK1B sequencing device (Oxford Nanopore Technologies). Base-calling was conducted with Guppy v3.2.10 on MinKnow 3.6.17 using a fast base-calling configuration. For Illumina MiSeq sequencing, DNA libraries were prepared using the Nextera XT library preparation kit and sequenced using a 2 X 250 bp paired-end configuration. The Illumina reads were trimmed using Trimmomatic v0.39 [14] and used for error correction using Pilon v1.23 [15]. For the genome assembly, multiple long-read assemblers were used: Flye v2.6 [16], Raven v1.1.10 (https://github.com/lbcb-sci/raven) and Miniasm-Minipolish v0.3 (https://github.com/rrwick/Minipolish). Then, Trycycler (https://github.com/rrwick/Trycycler) was used to obtain the consensus long-read assemblies from different long-read assemblers. The long reads were also used for error correction using Medaka v1.0.3 (https://github.com/nanoporetech/medaka). The genome was annotated using PGAP [17]. Additionally, antimicrobial resistance (AMR) genes were annotated using the Comprehensive Antibiotic Resistance Database (CARD) [18]. The genome and plasmids in UKMCC1015 were constructed using BRIG [19].

### Antimicrobial susceptibility testing and bacterial growth under different concentrations of NOR

The antimicrobial susceptibility profile of the UKMCC1015 strain was determined by the agar disk-diffusion method. The results were interpreted according to the CLSI guidelines. The NOR MIC for UKMCC1015 strains was determined using the broth-microdilution method. Escherichia coli 25922 was used as a reference control. The growth curve of UKMCC1015 was determined using 0X, 0.2X, 0.5X and 1.0X NOR MIC. The overnight culture of UKMCC1015 was diluted to an initial OD of 0.05 in 50 ml of broth containing the different antibiotic concentrations. The OD 600 nm was obtained every 15 minutes for 5.5 hours.

### Bacterial growth and RNA extraction

To study the short-term effect of a low dose of NOR on the transcriptome of UKMCC1015, overnight culture was diluted and allowed to grow in the presence of 0.2 X MIC NOR (37°C with shaking at 200 rpm) until the mid-log phase (OD 600 nm of 0.50) is reached. Cultures grown in the absence of norfloxacin were used as controls. RNA was extracted from the bacterial cells using the TRIzol phenol-chloroform extraction method according to the manufacturer’s protocol. rRNA depletion was performed using the MICROBExpress Bacterial mRNA Enrichment kit (Thermo Fisher Scientific, Massachusetts, USA) following the manufacturer’s protocol and cDNA library preparation using the NEBNext Ultra RNA Library Prep Kit for Illumina (New England Biolabs, Massachusetts, USA). DNA library quality control was done by measuring concentration using the Qubit 2.0 fluorometer (Invitrogen, California, USA) and assessing the size distribution using TapeStation 2200 (Agilent Technologies, California, USA). All libraries were pooled equimolar, and 15 pM was loaded into the MiSeq cartridge. Sequencing was performed on the MiSeq system using a 150 bp run configuration.

### RNA bioinformatics analyses

The *S. sonnei* UKMCC genome (GenBank assembly accession: GCA_014217935.1) was used as the reference genome to map the RNA sequencing data. Genome annotation was obtained from the NCBI PGAP annotation. *S. sonnei* Ss046 ncRNA sequences from the BSRD database [20] were used to BLAST against the *S. sonnei* genome to find additional ncRNA genes in the genome. BLAST hits were then appended to the annotation. To properly merge the two annotation sources, if annotations of the two sources overlapped, the PGAP annotation took precedence over the BLAST BSRD hits.

Illumina adapter sequences and low-quality bases were trimmed off using Trimmomatic v0.39. The trimmed, quality-filtered reads were mapped to the reference genome using Bowtie2 v2.3.5 and SAMtools v1.9 [21, 22]. The alignments on annotated genes were counted using htseq-count v0.11.2 [23]. Features included in the htseq counting were CDS, rRNA, ncRNA, tRNA, antisense RNA and riboswitch. Differential abundance analyses were conducted using DESeq2 [24] on the Galaxy platform, while the visualisation through the volcano plot was done using the web server VolcaNoseR [25]. The Clusters of Orthologous Genes (COGs) were determined using eggNOG (evolutionary genealogy of genes: Non-supervised Orthologous Groups), a web-based server [26]. Heatmaps were made using Python package seaborn [27] and matplotlib [28].

### Recombinant DNA techniques and oligonucleotides

Plasmid DNA and PCR products were extracted and purified using the FavorPrepTM Plasmid Extraction Mini Kit and the GEL/PCR Purification Kit according to the manufacturer’s instructions. For PCR, Q5® High-Fidelity, Taq DNA polymerases (New England Biolabs) and 2xTaq PCR MasterMix (SolarBio) were used according to the manufacturer’s recommendations. Cloning was performed using restriction enzymes (New England Biolabs & Vivantis Technologies), Quick CIP and T4 DNA ligase (New England Biolabs) described by the manufacturer. All primers used in this study were synthesised by Integrated DNA Technologies (Supplementary Table 2).

### Construction of ribosomal promoter-reporter plasmids

Plasmids were constructed using the restriction enzyme-based cloning approach. The promoter replacement approach was applied to clone different ribosomal protein (r-protein) promoters into the pUltra-RFP. R-protein promoter sequences of 151 nucleotides directly upstream of the transcriptional start site (TSS) were used in this analysis. The TSS was identified based on the TSS in *E. coli* [29]. The r-protein promoters rpsJp, rpsLp, rpsMp, rpsPp, rplNp, and rpsFp were amplified from UKMCC1015 gDNA and ligated to pUltra-RFP by EcoRI and PstI restriction sites. Plasmids constructed were transformed into E. coli DH10B via the heat-shock/calcium chloride procedure [30]. The constructs were verified by colony PCR and Sanger sequencing (Integrated DNA Technologies, IDT). The plasmids were extracted and transformed into UKMCC1015.

### Fluorescence assay

To analyse the effect of NOR on the promoter activity of various r-protein promoters, overnight cultures of the UKMCC1015 mutants were diluted to a starting OD of 600 nm of 0.05 and NOR, at a final concentration of 0.2X MIC, was added and was allowed to grow until it reaches the mid-log phase. Cultures without NOR stress served as controls. Then, the cells were spun down at 13,000 rpm for 5 minutes. The cell pellets were resuspended in 1X phosphate-buffered saline (1X PBS) of equal volume and vortexed. Then, 200 µL of the cells were added to a transparent, flat 96-well plate with a clear bottom. Relative fluorescence unit (RFU) was measured using the Infinite M200 (Tecan) microplate reader. Absorbance was measured at OD 600 nm, and the fluorescence signals were detected using 535 nm/620 nm (excitation/emission) and collected as top readings. Absorbance and fluorescence readings were corrected for background absorbance and fluorescence from the medium. The RFU value was normalised against the bacterial cell density (RFU/OD 600 nm) using the formula:

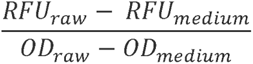

### Construction of UKMCC1015 Δ*relA* knockout and *relA* overexpression mutant

The knockout mutant (UKMCC1015 Δ*relA*) was constructed using homologous recombination. The wildtype UKMCC1015 (WT) was transformed with pGETrec plasmid, which carries the homologous recombination system, and selected using 100 µg/mL of AMP. DNA fragments with 300-bp homology arm upstream and downstream of tolC were amplified. The chloramphenicol-resistant (CmR) cassette was amplified from pBELOBAC11 plasmid. These 3 PCR fragments were subjected to overlap extension PCR to generate the Δ*relA* homologous template. The template was gel-purified using Monarch PCR & DNA Cleanup Kit (New England Biolabs, Massachusettes, USA) and transformed into UKMCC1015 electrocompetent cells. 1 mL of SOC medium was immediately added to the electroporated and incubated for 1 hour at 37°C with shaking before plating the culture on CHL agar to select for recombinants. UKMCC1015 Δ*relA* was verified by colony PCR and Sanger Sequencing. The pGETrec plasmid was removed from UKMCC1015 Δ*relA* by growing the strain for 40 generations with 0.2% L-arabinose and without AMP. To construct the overexpression mutant (UKMCC1015 *relA*^+^), *relA* was amplified from the UKMCC1015 gDNA and ligated to the pBAD24 vector, an arabinose-inducible plasmid, resulting in the construction of pBAD24_*relA.* This plasmid was transformed into *E. coli* DH10β and recovered on AMP agar. The plasmid was transformed into the UKMCC1015 after the construct was verified by colony PCR and Sanger sequencing (Integrated DNA Technologies, IDT).

### Laboratory evolution of NOR resistance

Overnight cultures of UKMCC1015, UKMCC1015 *relA^+,^* and UKMCC1015 Δ*relA* were grown in MHB broth at 37°C with 200 rpm shaking. Bacteria were subcultured onto fresh MHB broth in the presence of 1XMIC NOR daily for 14 days. The culture was then plated on MH agar containing different concentrations of NOR. Several colonies were selected from the NOR agar plates, and the new MIC of these colonies were determined using the broth microdilution method.

### Short-term NOR exposure

Overnight culture of UKMCC1015 evolved strains were diluted to an initial OD of 0.05 and grown to mid-logarithmic phase (OD 0.5) at 37°C with shaking at 200 rpm. For UKMCC1015 *relA*^+^, 0.2% of L-arabinose was added to the culture medium. At OD 0.5, the bacteria grew for another 2 hours in the presence of 1X MIC NOR. Then, RNA was extracted as described earlier.

### Determination of *relA* expression through semi-quantitative RT-PCR

Extracted RNA was treated with DNaseI and purified using Monarch RNA Cleanup Kit (New England Biolabs, Massachusetts, USA). The cDNA was synthesised from approximately 1000 ng of RNA using ReverTra Ace qPCR RT Master Mix with gDNA remover (TOYOBO, Japan) following the manufacturer’s protocol. Approximately 100 ng of cDNA for each sample was used for *relA* gene expression (using 16S rRNA as a housekeeping gene). The *relA* (4 µL) and 16S (2 µL) PCR products were run on 2% TBE agarose gel and post-stained with ViSafe Red Gel Stain (Vivantis Technologies) and visualised using GelDoc (BioRad). The relative intensity of the bands was quantified using ImageJ, and the expression of *relA* was normalised to the 16S rRNA expression levels.

## 4. Results

### Whole genome sequencing of *S. sonnei* UKMCC1015 revealed the presence of antimicrobial resistance (AMR) genes

We characterised the antimicrobial susceptibility profile and the complete genome of UKMCC1015 from Malaysia and investigated the transcriptomic changes of this bacterium to a sublethal NOR concentration. UKMCC1015 clinical strain was resistant to erythromycin but susceptible to other antibiotics such as ciprofloxacin sulfamethoxazole-trimethoprim, ampicillin-sulbactam, cefotaxime, nalidixic acid, as determined from disk diffusion assay (Fig. 1a). This strain was also sensitive to NOR (MIC of NOR = 0.03 µg/ml using the broth dilution method). Next, we investigated the presence of any AMR genes within the genome through whole-genome sequencing. A closed chromosome and two plasmids were obtained using hybrid assembly for UKMCC1015 (Fig. 1b, assembly information in Supplementary Table 3). The chromosome has a size of 4,926,332 bp, while the two plasmids, UKMCC1015_2 and UKMCC1015_3, have sizes of 7,231 and 2,101 bp, respectively, with an average GC content of 51.03% for the complete genome. 45 AMR genes associated with resistance to fluoroquinolone, macrolides, and tetracycline, among many other antibiotics, were identified from the UKMCC1015 genome (Fig. 1b, Supplementary Table 4). Most of the AMR genes (82%: 37/45 genes) encode for efflux pumps, while the others are responsible for antibiotic inactivation (4 genes) and drug target alteration (4 genes). 19 genes encoding for efflux pumps can potentially confer fluoroquinolone resistance. Besides, no mutations were observed in genes commonly associated with fluoroquinolone resistance: DNA Gyrase A and B (*gyrA* and *gyrB*) and DNA topoisomerase IV (*parC* and *parE*).

**Fig. 1:**
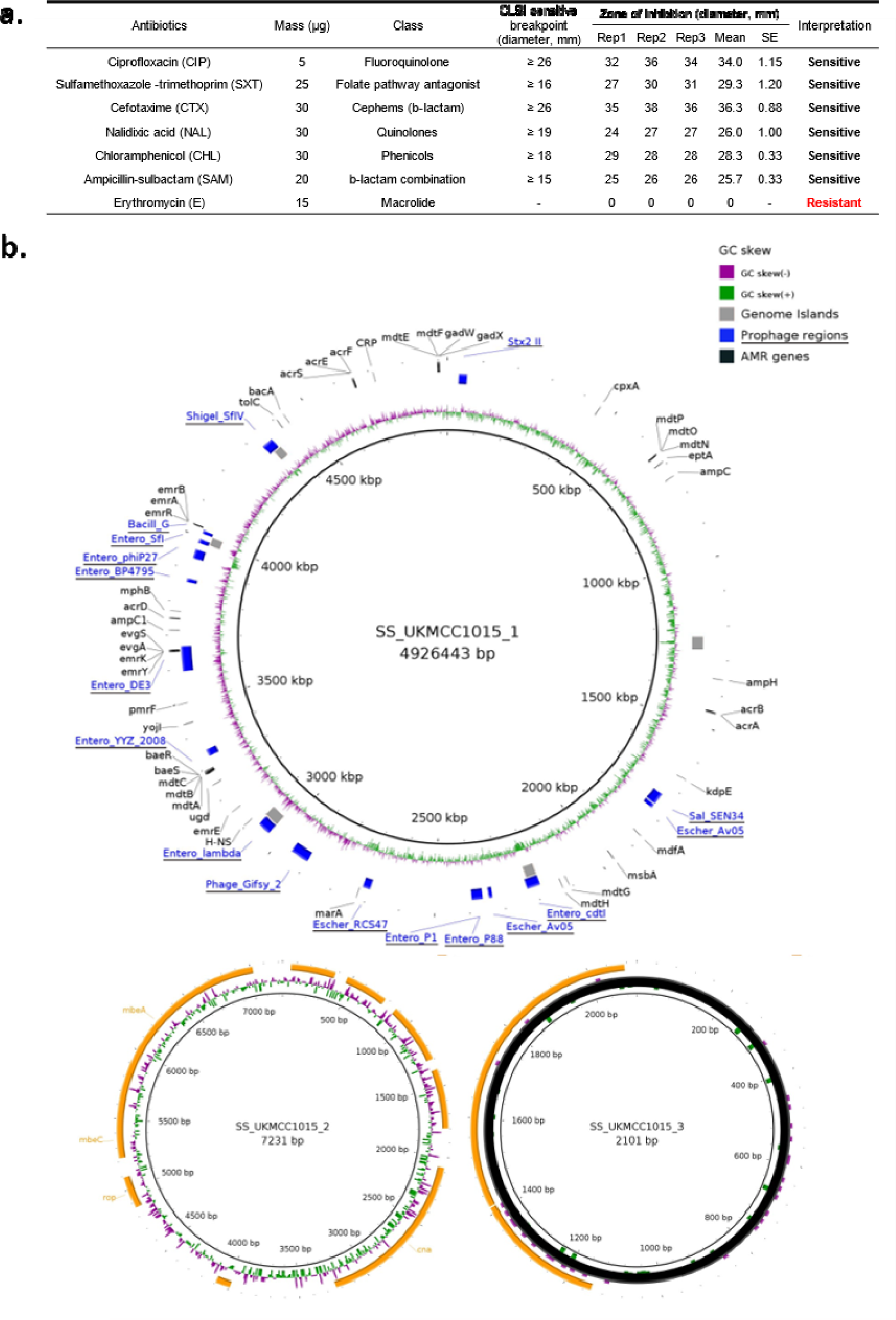
Phenotypic and genotypic characteristics of *Shigella sonnei* UKMCC1015. **a.** Antibiotic susceptibility profile of *S. sonnei* UKMCC1015. **b.** The closed chromosome of UKMCC1015, with UKMCC1015_1 (chromosome) and two plasmids, UKMCC1015_2 (bottom left) and UKMCC1015_3 (bottom right). UKMCC1015_1 chromosome, from inner ring outwards: GC skew, Genome Islands (Grey, obtained from IslandViewer predicted with at least 2 prediction methods), Prophage regions (Blue, underlined, obtained from Phaster), AMR genes (Black, obtained from Abricate using databases CARD). Plasmids of UKMCC1015, from the inner ring outwards: GC content, GC skew, genes in orange.

### Downregulation of translation in response to sub-lethal concentration of NOR

Based on the growth curve of UKMCC1015, 0.2X MIC NOR had the least effect on bacterial growth (Fig. 2a); hence, this sublethal NOR concentration was used in the subsequent RNA seq analysis. The total RNA UKMCC1015 grown on normal broth (control) and under 0.2X MIC NOR (experimental) was extracted, sequenced and analysed (Supplementary Table 5). The control samples were distinct from the experimental samples from the PCA plot (Supplementary Fig. 1). A total of 112 DEGs were identified in response to the sublethal NOR challenge (adjusted p-value ≤ 0.05, log_2_ (fold-change) ≥ 2). 31 genes show >2-fold upregulation, while 81 genes show >2-fold downregulation (Fig. 2b, Supplementary Table 6). Among these 112 genes, a non-coding RNA, *gcvB,* was differentially expressed. The remaining 111 genes were categorised using the COG database to elucidate further their role in UKMCC1015 response towards 0.2X MIC NOR (Fig. 2c). The top 5 categories were J (translation, ribosomal structure and biogenesis) with 49 genes, K (transcription) with 12 genes, S (function unknown) with 9 genes, F (nucleotide transport and metabolism) with 9 genes, and C (energy production and conversion) with 9 genes. None of the 45 AMR genes and the NOR targets found in UKMCC1015 were differentially expressed during 0.2X MIC NOR exposure (Fig. 2d).

**Fig. 2:**
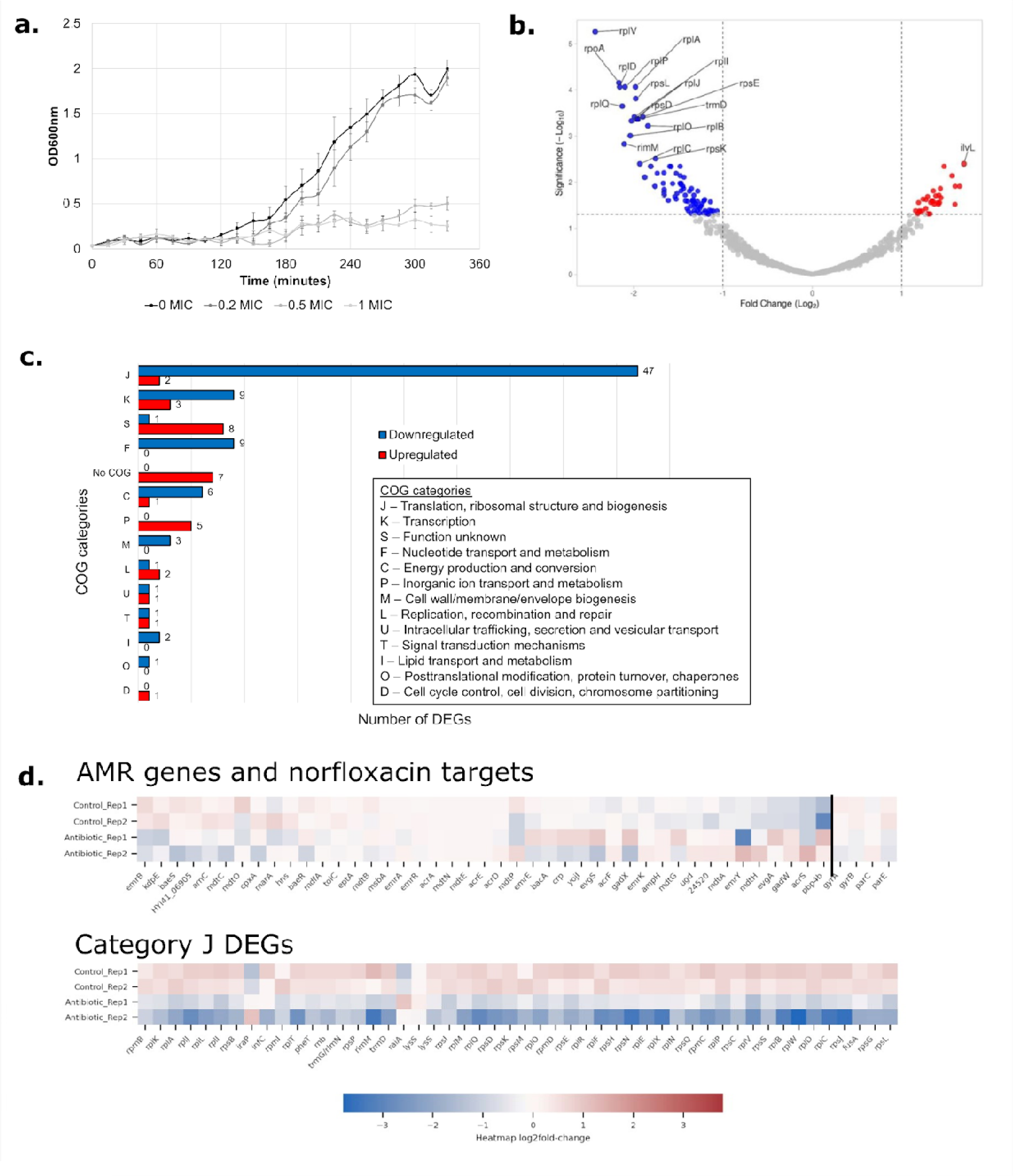
Effects of sub-MIC of norfloxacin towards UKMCC1015. **a.** Growth curve of UKMCC1015 WT under different sub-MIC of norfloxacin (0X, 0.2X, 0.5X and 1.0X MIC). **b.** Volcano plot of transcriptome analysis. **c.** COG categorisation of the 111 genes annotated by the non-supervised Orthologous Group (eggNOG). **d.** Expressions of all AMR genes and norfloxacin targets (*gyrA, gyrB, parC* and *parE*) which are not differentially expressed; and the expressions of the differentially expressed genes (DEGs) in Category J (translation, ribosomal structure and biogenesis).

Within category J, 47/49 DEGs were downregulated. The majority of the downregulated genes (37/47) encode ribosomal proteins (r-proteins), which are involved in the assembly of the ribosomal subunits (Shajani et al. 2011). Given that r-proteins are commonly expressed in operons, in which multiple genes are co-transcribed [31], we postulated that the downregulation of the genes in category J could be due to promoter regulation at the operon level. Indeed, 38/47 downregulated genes are expressed in 8 operons (Fig. 3a), supporting our idea that any regulation of these genes could be done through the promoters.

**Fig. 3:**
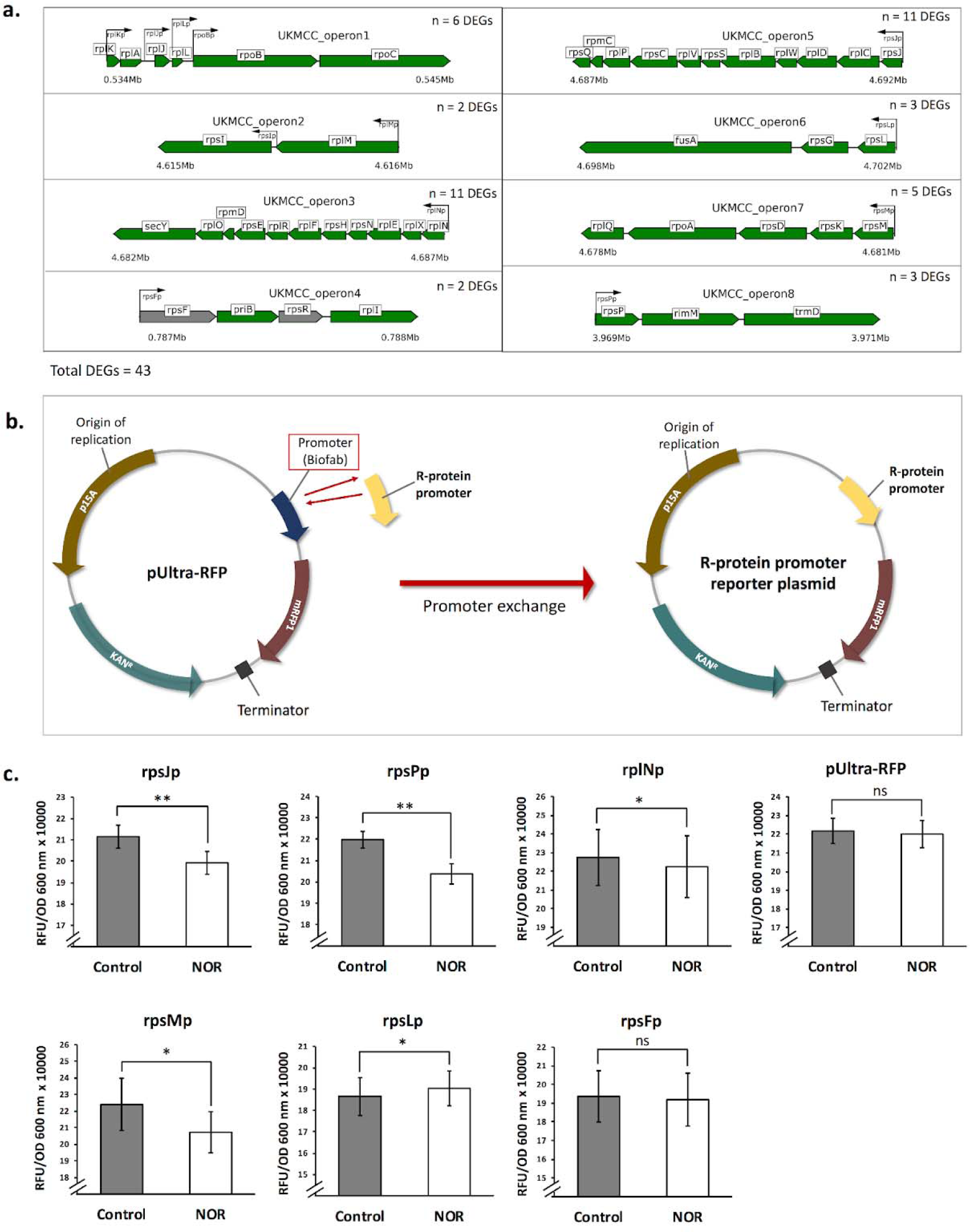
Schematic illustration of r-protein operons, promoter exchange approach and 0.2X MIC norfloxacin (NOR) effects on the promoter activity. **a.** Ribosomal protein operons of UKMCC1015. Green indicates differentially expressed genes (DEGs), and grey indicates non-differentially expressed genes. **b.** Construction of r-protein reporter plasmid via promoter exchange approach **c.** Normalised RFU/OD 600 nm changes for rpsJp, rpsMp, rpsPp, and rpsLp, rplNp, rpsFp r-protein promoters and pUltra-RFP promoter in response to 0.2X MIC norfloxacin stress. Three biological replicates were used for each promoter. Statistical significance (P-value < 0.05) was determined using Paired t-test (one-tailed, n=3); one asterisk (*): P-value < 0.05, two asterisks (**): P-value < 0.01, and ns: non-significant.

To further validate our hypothesis, promoter-reporter fusions were constructed (Fig. 3b), and the fluorescence readout from this reporter system was used to indirectly infer the activity of the r-protein promoters upon 0.2X MIC NOR exposure (Fig. 3c). Multiple promoters regulate the expression of operons 1 and 2. Therefore, these two operons are excluded from our analysis. Promoters from operons 3 - 8 were selected because these operons are under the control of only a single promoter (Fig. 3a), allowing direct measurements of the promoter activity on the operon transcription. The *rpsF* was not differentially expressed; hence, its promoter rpsFp was used as a control. The rpsJp, rpsPp, rpsMp, and rplNp promoters exhibited a slight decrease in the normalised RFU/OD 600 nm upon exposure to 0.2X MIC NOR with a percentage reduction of 5.81% (p-value = 0.0018), 7.28% (p-value = 0.004), 7.50% (p-value = 0.036), and 2.17% (p-value = 0.044), respectively, consistent with the RNA-seq findings, indicating that sub-lethal levels of NOR may reduce the activities of the r-protein promoters. The activity of the native promoter of the pUltra-RFP reporter plasmid that served as the negative control was also not significantly affected (Supplementary Fig. 2).

### ppGpp is not required for the evolution of NOR resistance

We hypothesised that sublethal NOR triggers the production of ppGpp and, in part, inhibits the expression of the r-proteins. To examine the contribution of ppGpp to the evolution of NOR resistance in UKMCC1015, we artificially overexpressed *relA* using the pBAD24 plasmid under the control of arabinose inducible promoter. Besides, we constructed a UKMCC1015 Δ*relA* mutant via homologous recombination to minimise the production of ppGpp. Overexpression of *relA* did not affect the intrinsic NOR resistance of UKMCC1015. However, the UKMCC1015 Δ*relA* strain had a NOR MIC 1.5x fold higher than the WT, indicating that UKMCC1015 is more resistant to NOR when the ppGpp production is minimal (Fig. 4a).

**Fig. 4:**
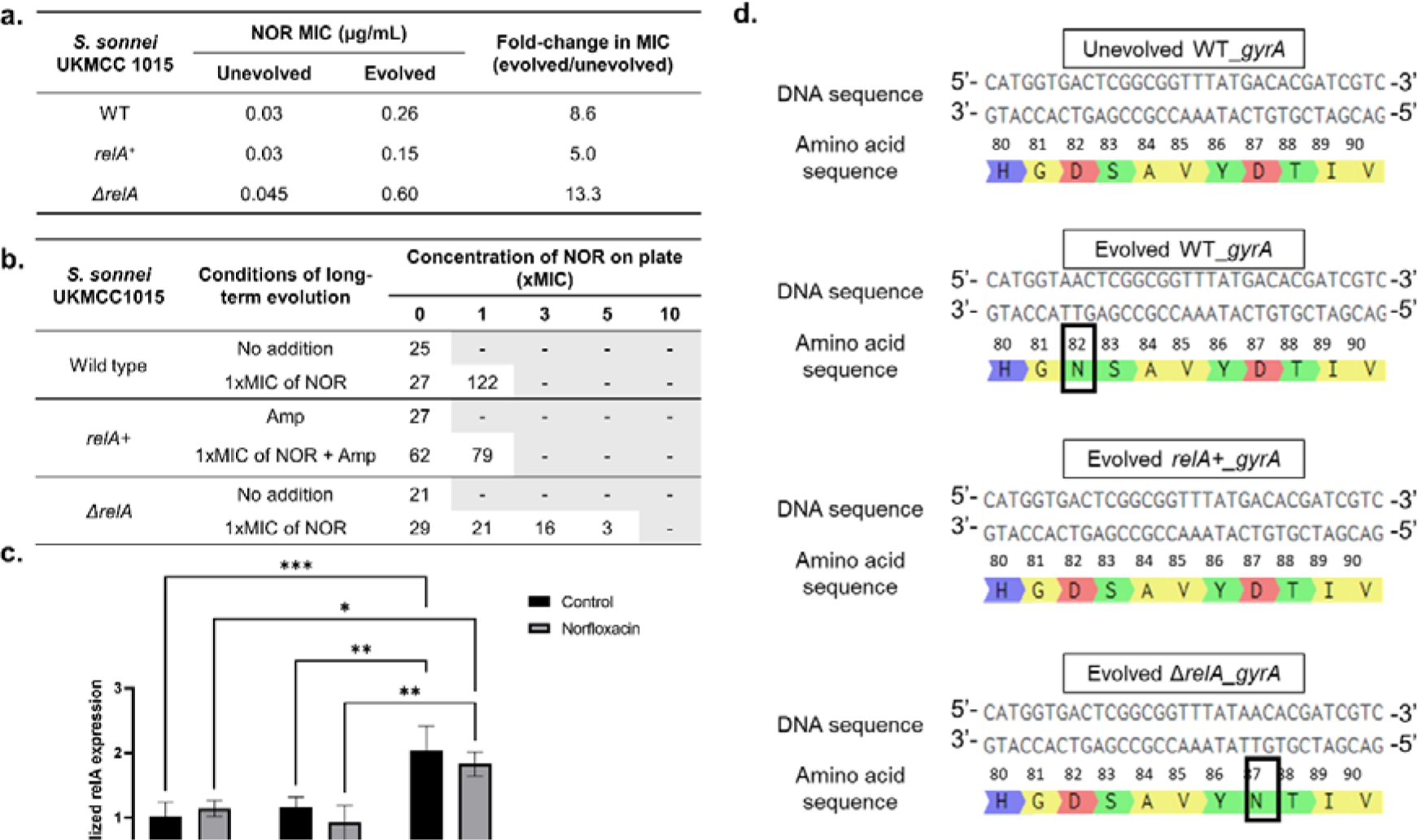
Response of UKMCC1015 and isogenic mutants to long-term exposure of NOR. **a.** MIC values towards NOR for unevolved and evolved strains. **b.** Number of colonies on agar containing different NOR concentrations after evolution. **c.** Semi-quantitative PCR results of *relA* and 16S rRNA before and after short-term exposure to NOR. Statistical significance (P-value < 0.05) was determined using two-way ANOVA; one asterisk (*): P-value < 0.05, two asterisks (**): P-value < 0.01, three asterisks (***): P-value < 0.001, four asterisks (****): P-value:<0.0001, and ns: non-significant. **d.** Mutations at amino acid position 80 - 90 in gyrA for unevolved WT, evolved WT, evolved *relA+,* and evolved Δ*relA*.

We evolved the UKMCC1015 *relA^+^*, UKMCC1015 Δ*relA* and UKMCC1015 WT (as a control) in 1.0X the respective NOR MIC for 14 days. The evolved cultures were plated on different concentrations of NOR. From the plating results (Fig. 4b), it can be seen that the Δ*relA* mutant grew at a concentration of 5xMIC, unlike the wild type and relA+ mutant, which can only grow up to 1xMIC. The evolved WT evolved UKMCC1015 *relA^+^* and evolved UKMCC1015 Δ*relA* had NOR MIC of 8.6-fold, 5-fold and 13.3-fold higher than their respective unevolved parental strain (Fig. 4a). This contradicts previous hypotheses where the increase of ppGpp levels are associated with the development of NOR resistance. We also observed that the evolved UKMCC1015 WT and evolved UKMCC1015Δ*relA* had single nucleotide polymorphism (SNP) mutations in the genes encoding for DNA gyrase A (Fig. 4d). We identified D82N and D87N in *gyrA* in our evolved UKMCC1015 WT and evolved UKMCC1015 Δ*relA* mutant, respectively. No SNP was identified in the *gyrA* of the evolved UKMCC1015 *relA^+^*.

Thus, we hypothesise that while ppGpp is needed during short-term exposure to NOR (around 2 hours), the prolonged expression of ppGpp under long-term NOR stress is counterproductive to the evolution of NOR resistance. To understand the role of ppGpp during short and long-term NOR stress, the expression level of *relA* in the evolved strains was further quantified using semi-quantitative RT-PCR (Fig. 4c, Supplementary Figure 3). The *relA* expression in evolved UKMCC1015 WT remains unchanged compared to unevolved WT. Overexpression of *relA* showed an approximately 2-fold increase in *relA* expression relative to WT, whereas UKMCC1015 Δ*relA* did not express *relA*, as measured by semi-quantitative RT-qPCR. For short-term exposure to 1X MIC NOR, no significant changes were observed in the *relA* expression levels for all strains relative to the no-antibiotic control.

## 5. Discussion

This study investigated the effects of sub-lethal exposure to a fluoroquinolone antibiotic, NOR, on *Shigella sonnei*, a pathogen responsible for shigellosis. UKMCC1015 was sensitive to most antibiotics tested, except for erythromycin, a macrolide antibiotic. As fluoroquinolones are the recommended treatment for shigellosis, norfloxacin was chosen for this investigation. The results showed UKMCC1015 (1) maintained the expressions of AMR genes and the NOR targets (DNA gyrase and topoisomerase IV) and (2) downregulated the expression of the r-proteins leading to reductions of translation, transcription, energy production and nucleotide production to cope with the sub-lethal NOR challenge.

Several past studies on fluoroquinolone-resistant *Shigella* indicate that the overexpression of genes encoding efflux pumps contributes to their resistance to fluoroquinolones [32–34]. Other responses towards fluoroquinolones include the downregulation of membrane porins to reduce membrane permeability and prevent antibiotic accumulation within the cell [35]. This suggests that the initial transcriptional response of fluoroquinolone-sensitive bacteria (UKMCC1015) is different from resistant ones. Instead of overexpressing the efflux pumps, the sensitive strain perhaps uses other pathways to cope with 0.2X MIC NOR. Additionally, there are no changes in the expressions of the NOR targets (DNA gyrase and topoisomerase IV). This may be due to the main fluoroquinolone resistance mechanism being linked to chromosomal mutations of *gyrA* and *parC* [35], which is not seen in UKMCC1015.

The transcriptomic analysis showed that exposure to sublethal NOR resulted in the under-expression of multiple r-protein operons even though r-proteins are not the NOR target. This observation was further supported using the promoter fluorescence assay. However, we observed only a slight reduction in the r-protein promoter activity compared to a larger reduction from the RNA-Seq analysis, suggesting that a decrease in the r-protein promoter activities could partly drive the repression of multiple r-protein gene transcripts. Besides, the rpsLp promoter activity was significantly upregulated by 2.08%. This discrepancy could be due to the delay between transcription and maturation of the fluorescent protein [36]. Overall, the large-scale downregulation of ribosomal protein operons suggests that a common regulator may modulate the response of UKMCC1015 to sublethal NOR exposure.

One such regulator is the alarmone guanosine tetraphosphate and guanosine pentaphosphate alarmone, collectively referred to as (p)ppGpp (ppGpp from here on). High ppGpp levels lead to the downregulation of multiple r-protein operons (Lemke *et al.* 2011). ppGpp is involved in the stringent response triggered during amino acid or fatty acid starvation or other environmental stress such as osmotic shock, heat shock and oxygen variation [37]. Some roles of ppGpp include growth rate control by inhibiting the production of ribosomal RNA and the regulation of general metabolism through the expression of genes in amino acid biosynthesis [38]. Besides, ppGpp may mediate the downregulation of nucleotide synthesis and, subsequently, a decrease in DNA replication. As a result, the bacteria decrease the effect of NOR on the DNA gyrase and topoisomerase IV, enzymes involved in DNA replication. This is also supported by how *gyrA* and *parC,* the targets of NOR, are not upregulated during sub-lethal NOR treatment. Instead of increasing the number of targets and DNA replication to overwhelm the concentration of antibiotics, as seen in another study [39], this fluoroquinolone-sensitive UKMCC1015 decreases its activities and becomes semi-dormant. Therefore, in UKMCC1015, the ppGpp level may be elevated in response to short-term sublethal NOR exposure, partly blocking the transcription of multiple r-proteins operons. However, alternative regulatory mechanisms controlling r-protein expressions cannot be ruled out.

Past studies showed that high levels of ppGpp help in antibiotic tolerance [40]. Hence, we investigated the role of ppGpp in modulating the adaptation of UKMCC1015 in response to long-term lethal NOR exposure. ppGpp is synthesised/hydrolysed by the RelA/SpoT homologues (RSH) superfamily proteins. The gene *relA* encodes for ppGpp synthetase with a single function, while *spoT* encodes for a bifunctional enzyme that mainly hydrolyses ppGpp with weak synthetase activity synthesis [41]. Given that the levels of ppGpp can be manipulated by overexpressing or deleting *relA*, the overexpression UKMCC1015 *relA+* and knockout UKMCC1015 Δ*relA* mutants were constructed to investigate the role of ppGpp in the long-term evolution of NOR resistance.

Interestingly, we observed that UKMCC1015 Δ*relA* has a slightly higher NOR MIC than the WT. The NOR MIC for UKMCC1015 *relA+*, on the other hand, is similar to the WT. These findings indicate that without *relA*, UKMCC1015 developed a slightly higher intrinsic resistance to NOR, similar to a study that found an increase in ampicillin tolerance in an *E. coli* Δ*relA* mutant in a minimal medium containing glucose after a prolonged stationary phase [42]. Others, however, have shown that knocking out *relA* leads to the loss of tolerance to ampicillin [43, 44], mecillinam [45], and microcin J25 [46]. This illustrates that ppGpp and stringent response may alter several aspects of antibiotic tolerance that lead to different tolerance levels.

Multiple studies reported the association between ppGpp and antibiotic tolerance/resistance. However, most were conducted quickly (within 48 hours) [44, 46–48]. We further showed that the evolution of UKMCC1015 WT and its isogenic mutants (UKMCC1015 *relA+* and UKMCC1015Δ*relA*) upon exposure to 1X MIC NOR (14 days) led to NOR resistance mainly via mutation in *gyrA* irrespective of ppGpp level. The intrinsic *relA* expression levels remain roughly constant between the unevolved and evolved strains, suggesting that ppGpp does not promote the long-term evolution of UKMCC1015 in this experimental condition. Instead, the specific mutation in the *gyrA* likely affects the degree of NOR resistance. The absence of *relA* led to a higher NOR MIC in the evolved UKMCC1015 Δ*relA*, approximately 2.3-fold higher than the evolved WT. This UKMCC1015 Δ*relA* developed a single mutation at position 87 (D87N), and this single nucleotide polymorphism (SNP) has been previously associated with fluoroquinolone resistance [49–51]. In evolved WT, we identified a mutation D82N. This previously unknown mutation may contribute to the 8.6-fold increase in NOR MIC because the SNP is located near one of the quinolone resistance-determining regions (Ser83). High ppGpp levels activate stringent responses as a first-line defence to cope with stressful conditions such as antibiotic exposure. Our study, however, shows that without *relA* (minimal ppGpp production), *gyrA* mutation could be the main pathway driving the evolution of NOR resistance in UKMCC1015. Interestingly, no mutation occurs in the UKMCC1015 *relA+* overexpression strain, yet we observed a moderate increase in NOR resistance (5-fold). Given that stringent response slows down DNA replication, artificially activating and maintaining stringent response (through *relA* overexpression) may hinder the development of resistance mutations because stringent response slows down DNA replication [52]. Therefore, artificially upregulating the expression of *relA* in our study could activate other regulatory mechanisms, resulting in a moderate increase in the NOR MIC.

Our findings suggest that the constant elevation of relA expression (high ppGpp level) is counterproductive to developing NOR resistance. It is hypothesised that the increase in ppGpp due to NOR exposure first slows down bacteria growth and translation as a transient response. ppGpp promotes DNA double-strand break (DSB) repair by stabilising backtracked RNA polymerase at the site of DNA breakage [53]. This allows the bacteria to develop NOR tolerance within the first few hours of exposure. However, increased ppGpp levels suppress bacterial growth, which is detrimental to bacterial survival. We postulate that high levels of ppGpp are beneficial during the initial stage of NOR exposure to allow for NOR tolerance but are detrimental in the long term. Thus, after 14 days of evolution, the UKMCC1015 *relA+* strain exhibits a lower proportion of survival at greater NOR concentration and a lower MIC than UKMCC11015 WT. In contrast, with reduced ppGpp levels, the ΔrelA mutant survives better under long-term exposure to NOR, with a higher MIC than the WT.

## 6. Conclusion

Overall, our genome and the transcriptomic studies provide new insights into how a fluoroquinolone-sensitive *S. sonnei* UKMCC1015 responds to a sublethal norfloxacin challenge. The commonly reported norfloxacin targets (DNA gyrase and topoisomerase IV) and AMR genes were not differentially expressed. However, many ribosomal protein genes were downregulated, which may represent a consequence of the activation of ppGpp-associated stringent response. This study provides new insights into the multitude of responses *S. sonnei* can have to a low concentration of antibiotics, including the involvement of the stringent response and the alarmone ppGpp. Our study highlights that the antimicrobial response of a sensitive strain to a sublethal antibiotic challenge differs from that of a resistant strain. Additionally, a prolonged period of elevated ppGpp levels may negatively affect the NOR tolerance of the bacteria. Future studies should focus on the exact pathways involving the stringent response that can aid in discovering potential targets for novel antibiotics or anti-resistance agents.

## Acknowledgements

This work was supported by the Fundamental Research Grant Scheme (FRGS) from the Ministry of Higher Education Malaysia (FRGS/1/2020/STG03/MUSM/03/1) and the Monash University Malaysia High Impact Research Support Fund Award 2022 (STG000174). We also thank Associate Professor Kumaran Narayanan for providing the pBAD24 plasmid.

## Ethical Statement

Not applicable

## Funding Statement

Wong Bao Chi: Monash University Malaysia High Impact Research Support Fund Award 2022 (STG000174)

Soffi Law Kei Kei: No funding support

Muhammad Zarul Hanifah Md Zoqratt: No funding support

Qasim Ayub: No funding support

Tan Hock Siew: FRGS Ministry of Higher Education Malaysia

## Data Accessibility

The complete genome of *Shigella sonnei* UKMCC1015 and its plasmids have been deposited at GenBank under accession numbers CP060117, CP060118 and CP060119. RNA-sequencing data are accessible through BioSample accession number SAMN16871737.

## Authors’ Contributions

Wong Bao Chi: experimental design, acquisition and interpretation of data, writing – drafting, review & editing

Soffi Law Kei Kei: experimental design, acquisition and interpretation of data, writing – drafting, review & editing

Muhammad Zarul Hanifah Md Zoqratt: data analysis

Qasim Ayub: writing – review & editing

Tan Hock Siew: supervision, conceptualisation, writing – review & editing

## Supplementary Information

Supplementary Table 1: Bacterial strains and plasmids used in this study.

Supplementary Table 2: Primers used in this study. Restriction sites used for cloning are underlined.

Supplementary Table 3: Genome assembly information of S. sonnei UKMCC1015.

Supplementary Table 4: List of antimicrobial resistance (AMR) genes in the genome of UKMCC1015, identified by Abricate using the CARD database.

Supplementary Table 5: RNA sequencing results of all 4 experimental sample groups.

Supplementary Table 6: Differentially expressed genes and non-coding RNAs in Shigella sonnei UKMCC1015 in response to 0.2xMIC of norfloxacin.

Supplementary Fig. 1: Principle component analysis (PCA) plot of RNA sequencing data. Gene expression changes towards 0.2X MIC of norfloxacin were investigated between the control and experimental groups.

Supplementary Fig. 2: Average normalised RFU/OD 600 nm, standard error, percentage changes and p-value for a. rpsJp, b. rpsPp, c. rpsMp, d. rpsLp, e. rplNp, f. rpsFp r-protein promoters, g. pUltra-RFP promoter for control and 0.2X MIC norfloxacin (NOR) exposure, and h. strain 5c rpsJp during 0.2X MIC norfloxacin stress. Experiments were conducted in triplicates.

Supplementary Fig. 3: Representative gel image for semi-quantitative PCR of relA and 16S rRNA gene.

## References

1. Baker, S., The, H. C. 2018 Recent insights into *Shigella*. Curr Opin Infect Dis. 31, 449–454. (10.1097/QCO.0000000000000475)

2. Thompson, C. N., Duy, P. T., Baker, S. 2015 The rising dominance of *Shigella sonnei:* an intercontinental shift in etiology of bacillary dysentery. PLoS Negl Trop Dis. 9, e0003708. (10.1371/journal.pntd.0003708)

3. Holt, K. E., Baker, S., Weill, F. X., Holmes, E. C., Kitchen, A., Yu, J., Sangal, V., Brown, D. J., Coia, J. E., Kim, D. W., et al. 2012 *Shigella sonnei* genome sequencing and phylogenetic analysis indicate recent global dissemination from Europe. Nat Genet. 44, 1056–1059. (10.1038/ng.2369)

4. Banga Singh, K. K., Ojha, S. C., Deris, Z. Z., Rahman, R. A. 2011 A 9-year study of shigellosis in Northeast Malaysia: Antimicrobial susceptibility and shifting species dominance. Z. fur Gesundheitswiss. 19, 231–236. (10.1007/s10389-010-0384-0)

5. Koh, X. P., Chiou, C. S., Ajam, N., Watanabe, H., Ahmad, N., Thong, K. L. 2012 Characterization of *Shigella sonnei* in Malaysia, an increasingly prevalent etiologic agent of local shigellosis cases. BMC Infect. Dis. 12, 122. (10.1186/1471-2334-12-122)

6. González-Plaza, J. J., Blau, K., Milaković, M., Jurina, T., Smalla, K., Udiković-Kolić, N. 2019 Antibiotic-manufacturing sites are hot-spots for the release and spread of antibiotic resistance genes and mobile genetic elements in receiving aquatic environments. Environ. Int. 130, 104735. (10.1016/j.envint.2019.04.007)

7. Bhattacharya, G., Dey, D., Das, S., Banerjee, A. 2017 Exposure to sub-inhibitory concentrations of gentamicin, ciprofloxacin and cefotaxime induces multidrug resistance and reactive oxygen species generation in meticillin-sensitive *Staphylococcus aureus*. J. Med. Microbiol. 66, 762–769. (10.1099/jmm.0.000492)

8. Lee, S. J., Park, N. H., Mechesso, A. F., Lee, K. J., Park, S. C. 2017 The phenotypic and molecular resistance induced by a single-exposure to sub-mutant prevention concentration of marbofloxacin in *Salmonella* Typhimurium isolates from swine. Vet. Microbiol. 207, 29–35. (10.1016/j.vetmic.2017.05.026)

9. Gumus, D., Kalayci-Yuksek, F., Yoruk, E., Uz, G., Celik, E., Arslan, C., Aydin, E. M., Canli, C., Ang-Kucuker, M. 2018 Alterations of growth rate and gene expression levels of UPEC by antibiotics at sub-MIC. Folia Microbiol (Praha). 63, 451–457. (10.1007/s12223-017-0582-z)

10. Bernier, S., Surette, M. 2013 Concentration-dependent activity of antibiotics in natural environments. Front. Microbiol. 4, (10.3389/fmicb.2013.00020)

11. Li, Z., Hu, Y., Yang, Y., Lu, Z., Wang, Y. 2018 Antimicrobial resistance in livestock: antimicrobial peptides provide a new solution for a growing challenge. Anim. Front. 8, 21–29. (10.1093/af/vfy005)

12. Aldred, K. J., Kerns, R. J., Osheroff, N. 2014 Mechanism of quinolone action and resistance. Biochemistry. 53, 1565–1574. (10.1021/bi5000564)

13. Ranjbar, R., Farahani, A. 2019 *Shigella:* antibiotic-resistance mechanisms and new horizons for treatment. Infect Drug Resist. 12, 3137–3167. (10.2147/IDR.S219755)

14. Bolger, A. M., Lohse, M., Usadel, B. 2014 Trimmomatic: a flexible trimmer for Illumina sequence data. Bioinformatics. 30, 2114–2120. (10.1093/bioinformatics/btu170)

15. Walker, B. J., Abeel, T., Shea, T., Priest, M., Abouelliel, A., Sakthikumar, S., Cuomo, C. A., Zeng, Q., Wortman, J., Young, S. K., et al. 2014 Pilon: An integrated tool for comprehensive microbial variant detection and genome assembly improvement. PLOS ONE. 9, e112963. (10.1371/journal.pone.0112963)

16. Kolmogorov, M., Yuan, J., Lin, Y., Pevzner, P. A. 2019 Assembly of long, error-prone reads using repeat graphs. Nat. Biotechnol. 37, 540–546. (10.1038/s41587-019-0072-8)

17. Tatusova, T., DiCuccio, M., Badretdin, A., Chetvernin, V., Nawrocki, E. P., Zaslavsky, L., Lomsadze, A., Pruitt, K. D., Borodovsky, M., Ostell, J. 2016 NCBI prokaryotic genome annotation pipeline. Nucleic Acids Res. 44, 6614–6624. (10.1093/nar/gkw569)

18. Alcock, B. P., Raphenya, A. R., Lau, T. T. Y., Tsang, K. K., Bouchard, M., Edalatmand, A., Huynh, W., Nguyen, A. V., Cheng, A. A., Liu, S., et al. 2020 CARD 2020: antibiotic resistome surveillance with the comprehensive antibiotic resistance database. Nucleic Acids Res. 48, D517–D525. (10.1093/nar/gkz935)

19. Alikhan, N. F., Petty, N. K., Ben Zakour, N. L., Beatson, S. A. 2011 BLAST Ring Image Generator (BRIG): simple prokaryote genome comparisons. BMC Genom. 12, (10.1186/1471-2164-12-402)

20. Li, L., Huang, D., Cheung, M. K., Nong, W., Huang, Q., Kwan, H. S. 2013 BSRD: a repository for bacterial small regulatory RNA. Nucleic Acids Research. 41, D233–D238. (10.1093/nar/gks1264)

21. Langmead, B., Salzberg, S. L. 2012 Fast gapped-read alignment with Bowtie 2. Nat. Methods. 9, 357–359. (10.1038/nmeth.1923)

22. Li, H., Handsaker, B., Wysoker, A., Fennell, T., Ruan, J., Homer, N., Marth, G., Abecasis, G., Durbin, R., Subgroup, G. P. D. P. 2009 The Sequence Alignment/Map format and SAMtools. Bioinformatics. 25, 2078–2079. (10.1093/bioinformatics/btp352)

23. Anders, S., Pyl, P. T., Huber, W. 2015 HTSeq--a Python framework for high-throughput sequencing data. Bioinformatics. 31, 166–169. (10.1093/bioinformatics/btu638)

24. Love, M. I., Huber, W., Anders, S. 2014 Moderated estimation of fold change and dispersion for RNA-seq data with DESeq2. Genome Biol. 15, 550. (10.1186/s13059-014-0550-8)

25. Goedhart, J., Luijsterburg, M. S. 2020 VolcaNoseR is a web app for creating, exploring, labeling and sharing volcano plots. Sci. Rep. 10, 20560. (10.1038/s41598-020-76603-3)

26. Huerta-Cepas, J., Szklarczyk, D., Heller, D., Hernández-Plaza, A., Forslund, S. K., Cook, H., Mende, D. R., Letunic, I., Rattei, T., Jensen, Lars J., et al. 2018 eggNOG 5.0: a hierarchical, functionally and phylogenetically annotated orthology resource based on 5090 organisms and 2502 viruses. Nucleic Acids Res. 47, D309–D314. (10.1093/nar/gky1085)

27. Waskom, M. L. 2021 seaborn: statistical data visualisation. J. Open Source Softw. 6, 3021. (10.21105/joss.03021)

28. Hunter, J. D. 2007 Matplotlib: A 2D Graphics Environment. Comput. Sci. Eng. 9, 90–95. (10.1109/MCSE.2007.55)

29. Lemke, J. J., Sanchez-Vazquez, P., Burgos, H. L., Hedberg, G., Ross, W., Gourse, R. L. 2011 Direct regulation of *Escherichia coli* ribosomal protein promoters by the transcription factors ppGpp and DksA. Proc. Natl. Acad. Sci. U.S.A. 108, 5712–5717. (10.1073/pnas.1019383108)

30. Sambrook, J., Russell, D. W. 2006 Preparation and Transformation of Competent *E. coli* Using Calcium Chloride. Cold Spring Harb. Protoc. 2006, pdb.prot3932. (10.1101/pdb.prot3932)

31. Osbourn, A. E., Field, B. 2009 Operons. Cell. Mol. Life Sci. 66, 3755–3775. (10.1007/s00018-009-0114-3)

32. Kim, J. Y., Kim, S. H., Jeon, S. M., Park, M. S., Rhie, H. G., Lee, B. K. 2008 Resistance to fluoroquinolones by the combination of target site mutations and enhanced expression of genes for efflux pumps in *Shigella flexneri* and *Shigella sonnei* strains isolated in Korea. Clin. Microbiol. Infect. 14, 760–765. (10.1111/j.1469-0691.2008.02033.x)

33. Yang, H., Duan, G., Zhu, J., Lv, R., Xi, Y., Zhang, W., Fan, Q., Zhang, M. 2008 The AcrAB-TolC pump is involved in multidrug resistance in clinical *Shigella flexneri* isolates. Microb. Drug Resist. 14, 245–249. (10.1089/mdr.2008.0847)

34. Taneja, N., Mishra, A., Kumar, A., Verma, G., Sharma, M. 2015 Enhanced resistance to fluoroquinolones in laboratory-grown mutants & clinical isolates of *Shigella* due to synergism between efflux pump expression & mutations in quinolone resistance determining region. Indian J Med Res. 141, 81–89. (10.4103/0971-5916.154508)

35. Fàbrega, A., Madurga, S., Giralt, E., Vila, J. 2009 Mechanism of action of and resistance to quinolones. Microb. Biotechnol. 2, 40–61. (10.1111/j.1751-7915.2008.00063.x)

36. Balleza, E., Kim, J. M., Cluzel, P. 2018 Systematic characterisation of maturation time of fluorescent proteins in living cells. Nat. Methods. 15, 47–51. (10.1038/nmeth.4509)

37. Pacios, O., Blasco, L., Bleriot, I., Fernandez-Garcia, L., Ambroa, A., López, M., Bou, G., Cantón, R., Garcia-Contreras, R., Wood, T. K., et al. 2020 (p)ppGpp and Its Role in Bacterial Persistence: New Challenges. Antimicrob. Agents Chemother. 64, e01283–01220. (10.1128/AAC.01283-20)

38. Hauryliuk, V., Atkinson, G. C., Murakami, K. S., Tenson, T., Gerdes, K. 2015 Recent functional insights into the role of (p)ppGpp in bacterial physiology. Nat. Rev. Microbiol. 13, 298–309. (10.1038/nrmicro3448)

39. Heo, A., Jang, H.-J., Sung, J.-S., Park, W. 2014 Global Transcriptome and Physiological Responses of *Acinetobacter oleivorans* DR1 Exposed to Distinct Classes of Antibiotics. PLOS ONE. 9, e110215. (10.1371/journal.pone.0110215)

40. Hobbs, J. K., Boraston, A. B. 2019 (p)ppGpp and the Stringent Response: An Emerging Threat to Antibiotic Therapy. ACS Infectious Diseases. 5, 1505–1517. (10.1021/acsinfecdis.9b00204)

41. Sinha, A. K., Winther, K. S. 2021 The RelA hydrolase domain acts as a molecular switch for (p)ppGpp synthesis. *Commun*. Biol. 4, 434. (10.1038/s42003-021-01963-z)

42. Varik, V., Oliveira, S. R., Hauryliuk, V., Tenson, T. 2016 Composition of the outgrowth medium modulates wake-up kinetics and ampicillin sensitivity of stringent and relaxed *Escherichia coli*. Sci Rep. 6, 22308. (10.1038/srep22308)

43. Rodionov, D. G., Ishiguro, E. E. 1995 Direct correlation between overproduction of guanosine 3’, 5’-bis pyrophosphate (ppGpp) and penicillin tolerance in *Escherichia coli*. J Bacteriol. 177, 4224–4229. (10.1128/jb.177.15.4224-4229.1995)

44. Kudrin, P., Varik, V., Oliveira, S. R., Beljantseva, J., Del Peso Santos, T., Dzhygyr, I., Rejman, D., Cava, F., Tenson, T., Hauryliuk, V. 2017 Subinhibitory Concentrations of Bacteriostatic Antibiotics Induce relA-Dependent and relA-Independent Tolerance to β-Lactams. Antimicrob Agents Chemother. 61, (10.1128/aac.02173-16)

45. Vinella, D., D’Ari, R., Jaffé, A., Bouloc, P. 1992 Penicillin binding protein 2 is dispensable in *Escherichia coli* when ppGpp synthesis is induced. EMBO J. 11, 1493–1501. (10.1002/j.1460-2075.1992.tb05194.x)

46. Pomares, M. F., Vincent, P. A., Farías, R. N., Salomón, R. A. 2008 Protective action of ppGpp in microcin J25-sensitive strains. J Bacteriol. 190, 4328–4334. (10.1128/jb.00183-08)

47. Rodionov, D. G., Pisabarro, A. G., de Pedro, M. A., Kusser, W., Ishiguro, E. E. 1995 Beta-lactam-induced bacteriolysis of amino acid-deprived *Escherichia coli* is dependent on phospholipid synthesis. J Bacteriol. 177, 992–997. (10.1128/jb.177.4.992-997.1995)

48. Goodell, W., Tomasz, A. 1980 Alteration of *Escherichia coli* murein during amino acid starvation. J Bacteriol. 144, 1009–1016. (10.1128/jb.144.3.1009-1016.1980)

49. Johnning, A., Kristiansson, E., Fick, J., Weijdegård, B., Larsson, D. G. J. 2015 Resistance Mutations in *gyrA* and *parC* are Common in *Escherichia* Communities of both Fluoroquinolone-Polluted and Uncontaminated Aquatic Environments. Front. Microbiol. 6, (10.3389/fmicb.2015.01355)

50. Mirzaii, M., Jamshidi, S., Zamanzadeh, M., Marashifard, M., Malek Hosseini, S. A. A., Haeili, M., Jahanbin, F., Mansouri, F., Darban-Sarokhalil, D., Khoramrooz, S. S. 2018 Determination of *gyrA* and *parC* mutations and prevalence of plasmid-mediated quinolone resistance genes in *Escherichia coli* and *Klebsiella pneumoniae* isolated from patients with urinary tract infection in Iran. J. Glob. Antimicrob. Resist. 13, 197–200. (10.1016/j.jgar.2018.04.017)

51. Yu, X., Zhang, D., Song, Q. 2020 Profiles of *gyrA* Mutations and Plasmid-Mediated Quinolone Resistance Genes in *Shigella* Isolates with Different Levels of Fluoroquinolone Susceptibility. Infect Drug Resist. 13, 2285–2290. (10.2147/idr.S257877)

52. Kraemer, J. A., Sanderlin, A. G., Laub, M. T. 2019 The Stringent Response Inhibits DNA Replication Initiation in *E. coli* by Modulating Supercoiling of oriC. mBio. 10, (10.1128/mBio.01330-19)

53. Zhai, Y., Minnick, P. J., Pribis, J. P., Garcia-Villada, L., Hastings, P. J., Herman, C., Rosenberg, S. M. 2023 ppGpp and RNA-polymerase backtracking guide antibiotic-induced mutable gambler cells. Mol Cell. 83, 1298–1310.e1294. (10.1016/j.molcel.2023.03.003)

