## Supplementary figures and images for "Adaptation of a fluoroquinolone-sensitive *Shigella sonnei* to norfloxacin exposure"

### Supplementary Fig 1

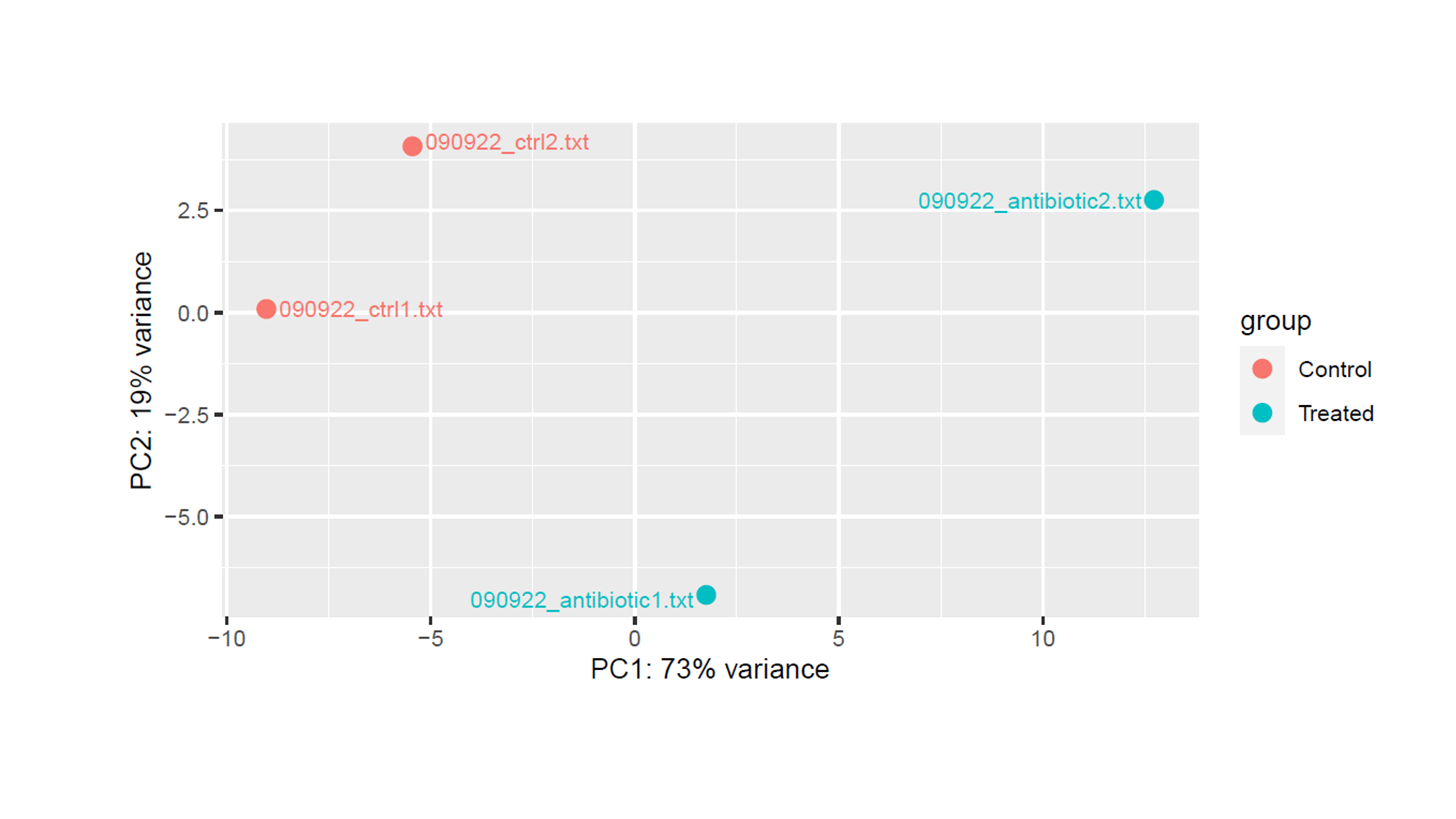

### Supplementary Fig 2

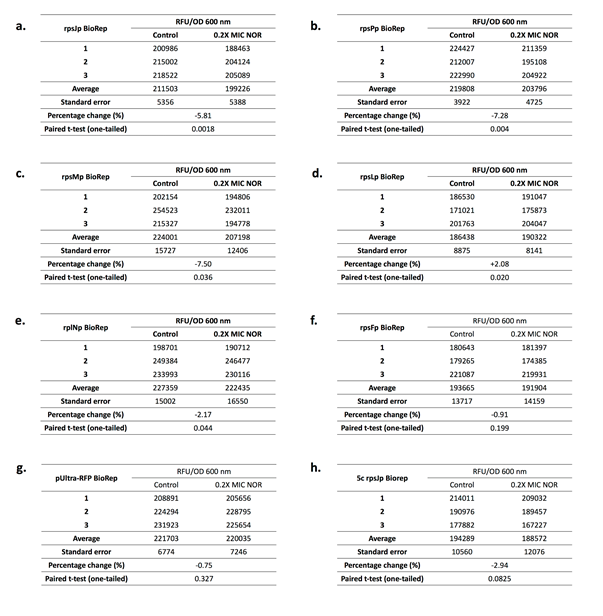

### Supplementary Fig 3

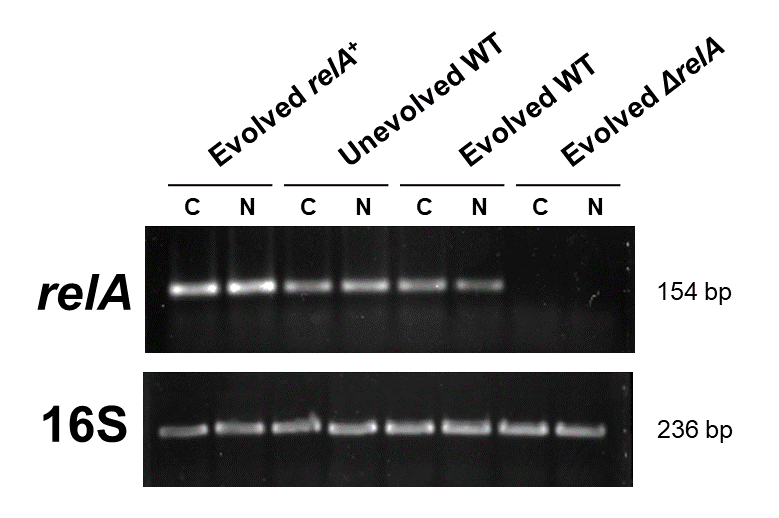
